# Neural Encoding for Human Visual Cortex with Deep Neural Networks Learning “What” and “Where”

**DOI:** 10.1101/861989

**Authors:** Haibao Wang, Lijie Huang, Changde Du, Dan Li, Bo Wang, Huiguang He

## Abstract

Neural encoding, a crucial aspect to understand human brain information processing system, aims to establish a quantitative relationship between the stimuli and the evoked brain activities. In the field of visual neuroscience, with the ability to explain how neurons in primary visual cortex work, population receptive field (pRF) models have enjoyed high popularity and made reliable progress in recent years. However, existing models rely on either the inflexible prior assumptions about pRF or the clumsy parameter estimation methods, severely limiting the expressiveness and interpretability. In this paper, we propose a novel neural encoding framework by learning “what” and “where” with deep neural networks. The modeling approach involves two separate aspects: the spatial characteristic (“where”) and feature selection (“what”) of neuron populations in visual cortex. Specifically, we use the receptive field estimation and multiple features regression to learn these two aspects respectively, which are implemented simultaneously in a deep neural network. The two forms of regularizations: sparsity and smoothness, are also adopted in our modeling approach, so that the receptive field can be estimated automatically without prior assumptions about shapes. Furthermore, an attempt is made to extend the voxel-wise modeling approach to multi-voxel joint encoding models, and we show that it is conducive to rescuing voxels with poor signal-to-noise characteristics. Extensive empirical results demonstrate that the method developed herein provides an effective strategy to establish neural encoding for human visual cortex, with the weaker prior constraints but the higher encoding performance.

**Author summary:** Characterizing the quantitative relationship between the stimuli and the evoked brain activities usually involves learning the spatial characteristic (“where”) and feature selection (“what”) of neuron populations. As an effective strategy, we propose a novel end-to-end “what” and “where” architecture to perform neural encoding. The proposed modeling approach consists of receptive field estimation and multiple features regression, which learns “where” and “what” simultaneously in a deep neural network. Different from previous methods, we use the sparsity and smoothness regularizations in the deep neural network to guide the receptive field estimation, so that the receptive field for each voxel can be estimated automatically. Moreover, in consideration of computational similarities between adjacent voxels, we made an attempt to extend the proposed modeling approach to multi-voxel joint encoding models, improving the encoding performance of voxels with poor signal-to-noise characteristics. Empirical evaluations show that the proposed method outperforms other baselines to achieve the state-of-the-art performance.

## Introduction

A great mystery in computational neuroscience is understanding how the brain effortlessly performs information perception and processing given sensory input. Uncovering this internal mechanism is of great scientific importance, not only for neuroscience researches, but also for artificial intelligence researches. In the field of visual neuroscience, one common method for insights into visual information processing is to establish neural encoding models with functional magnetic resonance imaging(fMRI) [1, 2]. This modeling approach (Fig 1) of neural encoding links fMRI signals at the millimeter scale to neural response at the micron scale, providing a non-invasive approach to revealing the nonlinear relationship between the external stimuli and the evoked brain activities [3–5].

**Fig 1.**
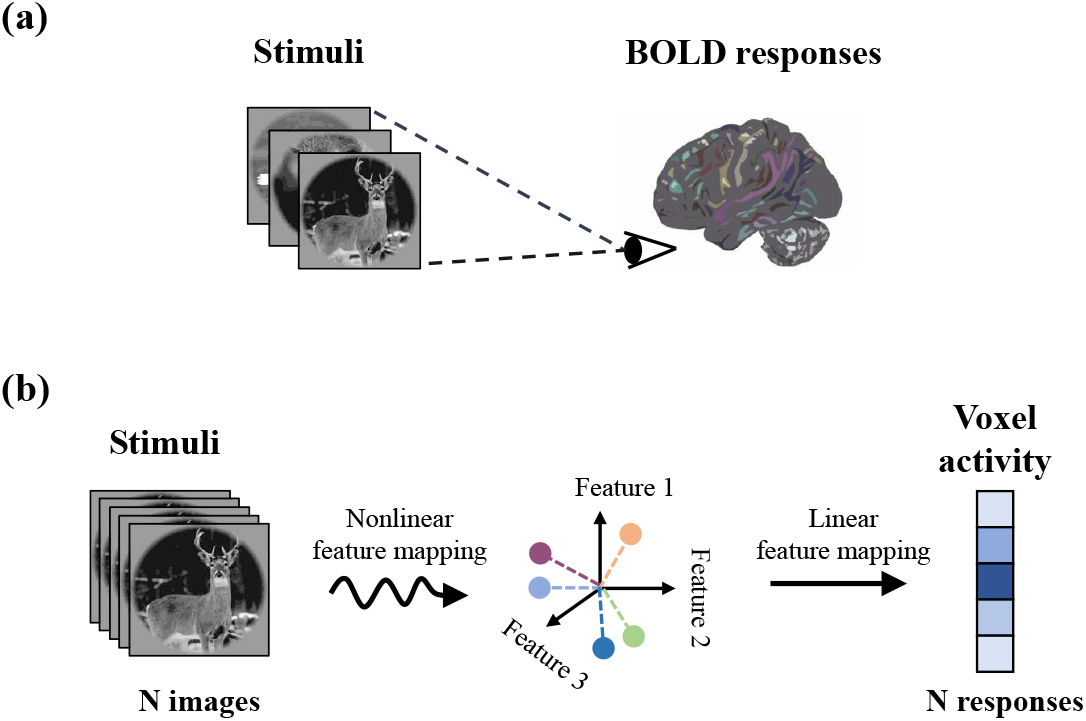
The general approach of neural encoding models. (a) Voxel activity evoked by experimental stimuli (such as natural images) is collected. (b) Encoding models are specified by a nonlinear mapping (curvy arrow) of the stimuli into an abstract feature space, as well as a linear mapping (straight arrow) from feature space to voxel activity.

Numerous studies have built neural encoding models from diverse perspectives for human visual cortex, including, but not limited to, Gabor wavelet pyramid model [4], luminance contrast models [6–10], the motion energy model [11], semantic models [12], deepnet models [13], and other machine learning methods [14, 15]. The estimated characteristics of visual cortex derived from these models can be roughly summarized in two aspects: “what” and “where”. “Where” characterizes the spatial characteristic of neuron populations in visual cortex, i.e. the location and extent of pooling over visual features, whereas “what” characterizes the feature selection property of neuron populations in visual cortex.

In general, “where” is based on the classical receptive fields. In the population receptive field (pRF) model [6], the visual feature is a binary map of the pixels occupied by a high-contrast stimulus (e.g., bar, ring, wedge). For each voxel, the model is constructed by an isotropic Gaussian area, the pRF, that pooled the visual feature map within a spatially localized area. This seminal approach quantitatively measured the population receptive field properties in human visual areas for the first time and was extended to some similar pRF models [7–10]. “What” focuses on feature selection and feature tuning functions. In the semantic model [12], several object category features (e.g. the presence of an animal, or a car.) are encoded as the vector of binary variables. Then, every object category was assigned a tuning parameter for each voxel to construct the model. Similarly, different visual features are further studied in subsequent models [4, 11, 14–16].

Recently, deep learning with neural networks [17–19] has been widely used to perform feature learning from scratch with promising performance, which sparks interest in using deep learning methods for understanding information processing in visual cortex [5, 20–22]. Based on deep neural networks, [13] proposed a new approach to encoding visual features named feature-weighted receptive field (fwRF) [13]. It starts with a natural image, obtains feature maps in a pre-trained convolutional neural network, and computes a weighted sum within the spatial extent of an 2D Gaussian receptive field. Finally, it regresses all the feature maps onto brain activity simultaneously, which yielded the state-of-the-art prediction accuracy. However, while previous work demonstrated promising results of processing in visual cortex, neural encoding models still lack adequate examination and require plenty of effort to improve.

There are two main challenges getting in the way of development of effective models. On one hand, conventional approaches [6, 7] are endowed with inflexible prior assumptions on the spatial characteristics of receptive fields, which limit the effectiveness of models to a large extent. For example, in the population receptive model [6], it assumed that the pRF has an isotropic Gaussian topography while the potentially suppressive surround is neglected. There have been subsequent models [7, 9, 10, 13] which have adopted the same principles with different pRF topographies. In general, one assumption about receptive field structure puts one prior constraint on the ability to extract the receptive field topography of the model. Specifically, inaccurate assumptions about receptive field topography may lead to erroneous estimation of the receptive field. Hence, it is meaningful to propose a new method that can extract receptive field topography without inflexible prior assumptions. On the other hand, previous approaches [6, 7, 9, 10, 13, 15] to obtaining receptive fields are based on grid search, which set search parameters according to experience. Accordingly, they are prone to pRF center mislocalization and size miscalculation. In the population receptive models [6, 7, 13], the pRF topography parameters were set to certain parameters, which can be obtained by minimizing the residual between the observed fMRI signal and the predicted signal. In this case, according to different shape parameters (i.e. center and radius), these models will inevitably generate large quantities of candidate pRF. That is, the grid fitting requires searching over quite large model-parameter spaces. Consequently, their encoding performances depend on the amount and parameter interval of candidate receptive fileds, which are set artificially. Obviously, more often than not, it is not optimal and requires lots of manual effort. It would therefore be significant to obtain the receptive fileds automatically in a more reasonable way.

Existing methods are prone to suffer from one or both of these issues and yield dissatisfaction. Attaching great importance to these bottlenecks, we proposed a novel “what” and “where” neural encoding architecture via deep neural networks. The proposed method first extract hierarchical features from the DNN driven for image recognition. Then, the original features are refined via channel attention and spatial receptive field, and finally are regressed simultaneously onto voxel activity. Different from previous methods, we use the sparsity and smoothness regularizations in the deep neural network to guide the receptive field estimation. This modelling approach can estimate the receptive field for each voxel automatically and maintain powerful expressiveness. In consideration of computational similarities, we extend the voxel-wise modeling approach to multi-voxel joint encoding models, which is beneficial to rescuing voxels with poor signal-to-noise characteristics. Our main contributions can be summarized as follows.

- We provide a new perspective on the deep-learning-based neural encoding models, performing receptive field estimation and features regression simultaneously in a deep neural network. This modeling approach can yield explicit receptive fields (“where”) and feature turning functions (“what”) automatically, which is rich in interpretability.
- The estimation of receptive fields is endowed with weaker constraints. Instead of strong prior assumptions on the shape of receptive fields, L1 regularization and Laplacian smoothing are adopted in our modeling approach, which can be regarded as weak prior assumptions about receptive fields.
- We made an attempt in the extension of the modeling approach. In consideration of the computational similarities between voxels, the voxel-wise modeling approach is extended to multi-voxel joint encoding models, suggesting a new approach to rescuing voxels with poor signal-to-noise characteristics more effectively.
- Extensive empirical evaluations on the publicly available fMRI dataset demonstrate that our modeling approach achieves superior performance compared with other neural encoding models.

## Methods

In the neural encoding dataset, we assume **X** = [**x**_1_, …, **x**_*N*_]^**T**^ ∈ ℝ^*N* ×*M*^ and **Y** = [**y**_1_, …, **y**_*N*_]^**T**^ ∈ ℝ^*N ×D*^ denote the matrices of visual images and the evoked fMRI activities, respectively. Here, *N* denotes the size of the training set. *M* and *D* denote the dimensions of visual image and fMRI activity pattern, respectively. Given an image **x**_*i*_, its hierarchical visual features can be obstained from a pretrained deep neural network (e.g., AlexNet [18]). Here, **H** = [**h**_1_, …, **h**_*N*_]^**T**^ ∈ ℝ^*N ×K*^ donotes the intermediate DNN features, where *K* denotes the number of feature maps. For modeling the statistical relationship between the visual images and the evoked voxel activities, we put forward a novel neural encoding framework by learning “what” and “where” based on deep neural networks. The simplified illustration of the proposed modeling approach is shown in Fig 2. Formally, it consists of three cascaded stages: 1) nonlinear feature extraction: extracting hierarchical visual features through a pretrained DNN model. 2) nonlinear feature refinement: converting original features into refined features with channel attention module and spatial receptive field (RF) module. 3) voxel-wise linear mapping: regressing refined features simultaneously onto voxel activities. In the following, we present the proposed approach in detail.

**Fig 2.**
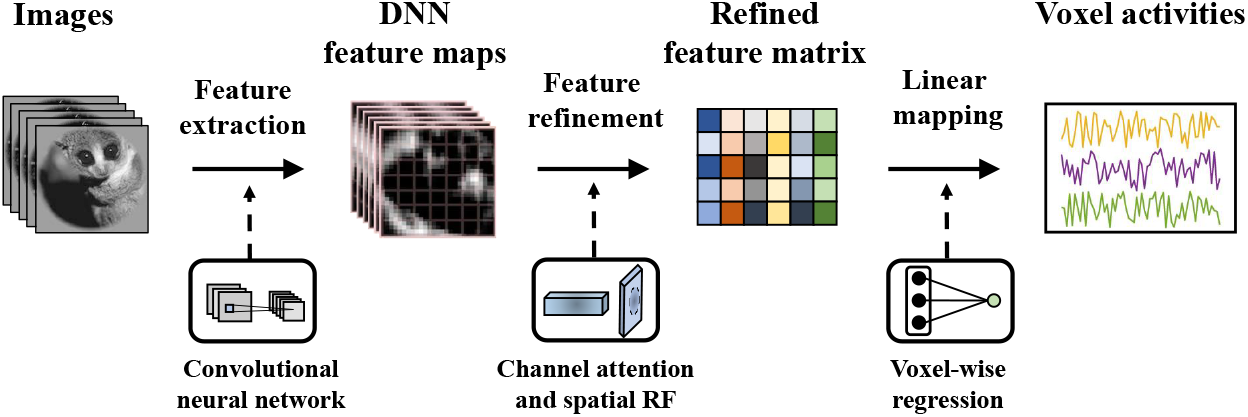
An overview of the proposed neural encoding framework. It consists of three cascaded stages: 1) nonlinear feature extraction: extracting hierarchical visual features through a pretrained DNN model. 2) nonlinear feature refinement: converting original features into refined features with channel attention module and spatial receptive field (RF) module. 3) voxel-wise linear mapping: regressing refined features simultaneously onto voxel activities.

### Nonlinear feature extraction

The neural encoding models take visual stimuli as the input, and output the evoked brain activities. Normally, it includs two sequential stpdf. The first step is a nonlinear feature mapping, converting the visual input to its feature representations; the second step is a voxel-wise linear mapping, projecting the feature representations onto activities at each voxel [4, 23–27]. In the present study, The nonlinear feature mapping consists of two parts: nonlinear feature extraction and nonlinear feature refinement. The nonlinear feature extraction is introduced in this sub-section, while nonlinear feature refinement and voxel-wise linear mapping are described in the next several sub-sections.

Recent studies [13, 28] has demonstrated that Alexnet [18], a specific version of DNNs, is capable of predicting voxel activities with statistical significance and high accuracies throughout the visual cortex. In line with previous work [13], a deep neural network (a specific implementation referred as the AlexNet) is adopted to extract nonlinear features in the present study. Briefly, AlexNet has been pre-trained to achieve the best preformance in Large Scale Visual Recognition Challenge 2012 [18]. It consists of eight layers of computational units: the first five layers were convolutional layers, while the rest layers were fully connected. The image input was fed into the first layer; the output from one layer served as the input to its next layer. Each convolutional layer involves plenty of units and a set of filters that extracts filtered outputs from different locations of the input. The resolution (square root of the number of pixels in each feature map) and depth (number of feature maps) for each convolutional layer was (55, 96), (27, 256), (13, 384), (13, 384), (13, 256) respectively. The fully-connected layers contained 4096, 4096 and 1000 units respectively. All feature maps from all convolutional layers, as well as up to 1024 units from the fully-connected layers, are used to result in hierarchical nonlinear features in the present study. Once trained, the CNN is able to undergo feature extraction by a simple feedforward pass of an input image. Herein, we refer to **h**_*i*_ as the intermediate DNN features given image **x**_*i*_. In consideration of feature maps from multiple layers may be adopted in the model, we index each layer by *l*, and let 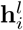 denotes the features from the *l*-th layer.

A question arises whether multiple feature maps included in the model are meaningful. In reality, we do not konw what features can better explain activity in the visual cortex under many circumstances. In this way, the total feature maps **h**_*i*_ are able to contain adequate features to capture the reasonable hypotheses about what is encoded in the visual cortex. Based on these original feature maps, the proposed method can infer which features and locations are important for explaining the activity in the voxel. In the nonlinear feature refinement, the feature map pixels are focused on within the spatial RF, as concentrating on important locations and suppressing unnecessary ones are conducive to improving the encoding performance. In this way, spatial RF is the “where” parameters of the neural encoding model. In the voxel-wise linear mapping, each feature map will be assigned an associated feature weight, which indicates the importance of the feature map for predicting the activity of each voxel. In this sense, all the feature map weights are “what” parameters of the neural encoding model. Noting that while original features maps are same for each voxel, but the feature refinement and feature weights vary across voxels.

### Nonlinear feature refinement

The crucial part of the proposed method is the nonlinear feature refinement, which contributes to encoding performance promotion and interpretability enhancement. In this subsection, we sequentially employ the channel attention module and spatial RF module whereby the proposed method can learn “where” to attend in visual information processing. The illustration of this feature refinement is shown in Fig 3.

**Fig 3.**
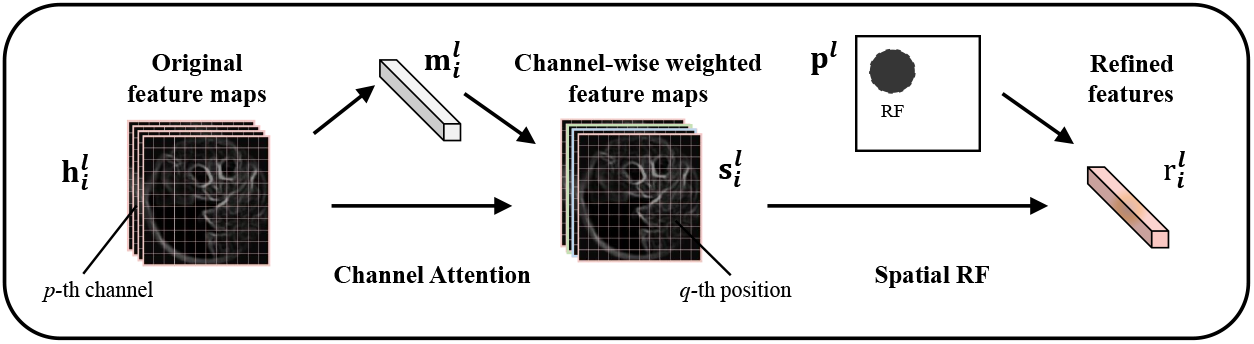
The feature refinement in the specific layer (the *l*-th layer). In the channel attention module, the original feature maps 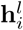 are initially used to obtain the channel attention weights 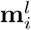. In the next phase, the original feature maps are element-wise multiplied by the channel attention weights to obtain the channel-wise weighted feature maps 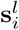. In the spatial RF module, the receptive field **p** is reshaped to the corresponding receptive field **p**_*l*_ according to the size of channel-wise weighted feature maps 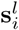. The Hadamard product of **p**_*l*_ and 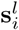 finally produce the refined features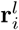.

### Channel attention module

It is acknowledged that attention plays an crucial role in human perception [29–31]. One important property of the human visual system is that one does not attempt to process a whole scene at once. Instead, humans exploit a sequence of partial glimpses and selectively focus on salient features in order to capture visual structure better [32–34]. In the present work, we pay more attention to the meaningful feature maps rather than considering each feature map equally. Since a channel-wise feature map is a detector response map of the corresponding filter in essence, channel attention can be regarded as the process of selecting inter-layer feature attributes and reducing redundant information (strengthening important ones and weakening unimportant ones).

Here, the channel attention module is built up based on the hierarchical features **h**_*i*_. Without loss of generality, we discard the image subscript *i* and layer-wise superscrip *l*. In each specific convolutional layer, the original feature maps 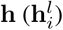 are reshaped to **h** = [**h**_1_, **h**_2_, …, **h**_*C*_] ∈ ℝ^*S*×*C*^, where **h**_*k*_ donotes the *k*-th channel of the feature maps. *C* and *S* are the channel dimension and spatial dimension in this layer, respectively. Given the input **h**, global average pooling is applied to each channel to obtain the mean vector 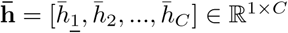 were 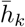 is the channel mean of **h**_*k*_. On the basis of the mean vector 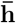, the channel attention module Φ(·) can be further formulated as follows:

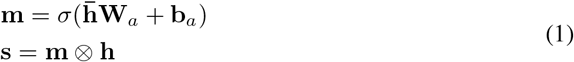

where **W**_*a*_ ∈ ℝ^*C*×*C*^ and **b**_*a*_ ∈ ℝ^1*×C*^ are the transformation matrix and bias term respectively; *σ* denotes the sigmoid function and ⊗ denotes element-wise product; **m** ∈ ℝ^1*×C*^ denotes the attention weight while **s** ∈ ℝ^*S×C*^ stands for attentioned feature maps. During element-wise multiplication, the channel attention weight is broadcasted (copied) along the spatial dimension. In this way, the original feature maps **h** are converted into attentioned feature maps **s**.

### Spatial RF module

In visual areas, population activity at each voxel in the cortical sheet encodes visual features within a spatially localized region of visual field [6–9]. In view of this, the attentioned feature maps are pooled within a limited and contiguous area in this module, which is the spatial receptive field (RF). In order to overcome the drawbacks brought by the inflexible prior assumptions or the clumsy parameter estimation methods, the sparsity regularization and smoothness regularization are adopted in this module. Under the guidance of both regularizations, the proposed method yields explicit receptive field automatically and encodes features within a contiguous region of the visual field.

Mathematically, the receptive field is denoted as **p** given each specific voxel, and randomly initialized with the same spatial size of natural images (227 × 227). In consideration of different sizes of feature maps, the receptive field is adapted to different convolutional layers by the means of reshaping. Taking the first convolutional layer as an example, the corresponding receptive field size is 55 × 55. In this way, there are five sizes of receptive fields converted from the original receptive field, with one-to-one correspondence to the five convolutional layers. As a direct way to combine attentioned feature maps with receptive fields, element-wise product is more natural and general than specially designed operations. Herein, the receptive field is broadcasted (copied) along the channel dimension in the beginning, matching with the dimensions of attentioned feature maps. Given the feature maps **s** = [**s**_1_, **s**_2_, …, **s**_*C*_] ∈ ℝ^*S×C*^, the spatial RF module Ψ(·) can be formulated as follows:

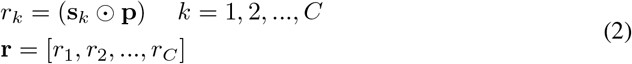

where **s**_*k*_ ∈ ℝ^*S*×1^ is the *c*-th channel of attentioned feature maps and **p** *∈* ℝ^*S×*1^ is the voxel-wise receptive field. ⊙ is the Hadamard product of matrices. Hence, spatial RF module Ψ(·) outputs the refined feature vector **r** ∈ ℝ^1×*C*^.

As a matter of fact, if there is no any other operation applied to the receptive field, it is just an ordinary mask. Inspired by the physiological structural characteristics of receptive field, our model adopted two forms of regularization: sparsity and smoothness. Specifically, for each specific voxel, since we expect its receptive field to be highly sparse, the receptive field was regularized by L1 penalty with strength *λ*_*s*_:

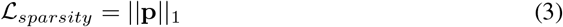

To ensure the receptive field focuses on an localized area as effectively as possible, we use an L2 penalty on the Laplacian of the receptive field with the strength *λ*_*l*_:

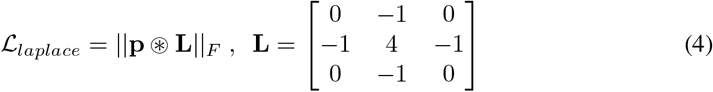

Here, ‖·‖_*F*_ and ‖·‖_1_ denote Frobenius norm and *L*_1_ norm (“entriwise” norm) of a matrix respectively, and ⊛ is the convolutional operation. In this way, sparsity and smoothness regularizations make the ordinary mask transform into the meaningful receptive field. In contrast to Gaussian assumption, these two forms of regularization can be viewed as weak prior assumptions about receptive field, which is more conducive to the expressiveness and flexibility of neural encoding models.

### Voxel-wise linear mapping

The original intention of neural encoding models is to account for the responses of different visual processing stages and reveal the information processing mechanism of neurons in visual cortex. Voxel-wise linear mapping sets up such a computational path to relate visual features to the evoked response at each voxel, bridging the gap between feature selection property of neuron populations in visual cortex and hierarchical feature maps in deep neural networks.

In some previous studies [21, 28, 35], the voxel-wise encoding models regress feature maps in each layer onto brain activity independently, which is proved highly effective. However, it reduces the model scale at the expense of model expressiveness. In reality, increasingly abstract and complex visual features are encoded in the deep neural networks (AlexNet). The convolutional layer encoded location and orientation-selective features, whereas the fully-connected layer encoded semantic features. In the light of feature diversity, we use feature maps from all layers to predict each voxel’s response rather than assume an one-to-one correspondence between a voxel and a DNN layer. It is a reasonable way that each feature map is assigned an appropriate weight, which can be learned from training data directly by the appropriate optimization algorithm.

**Fig 4.**
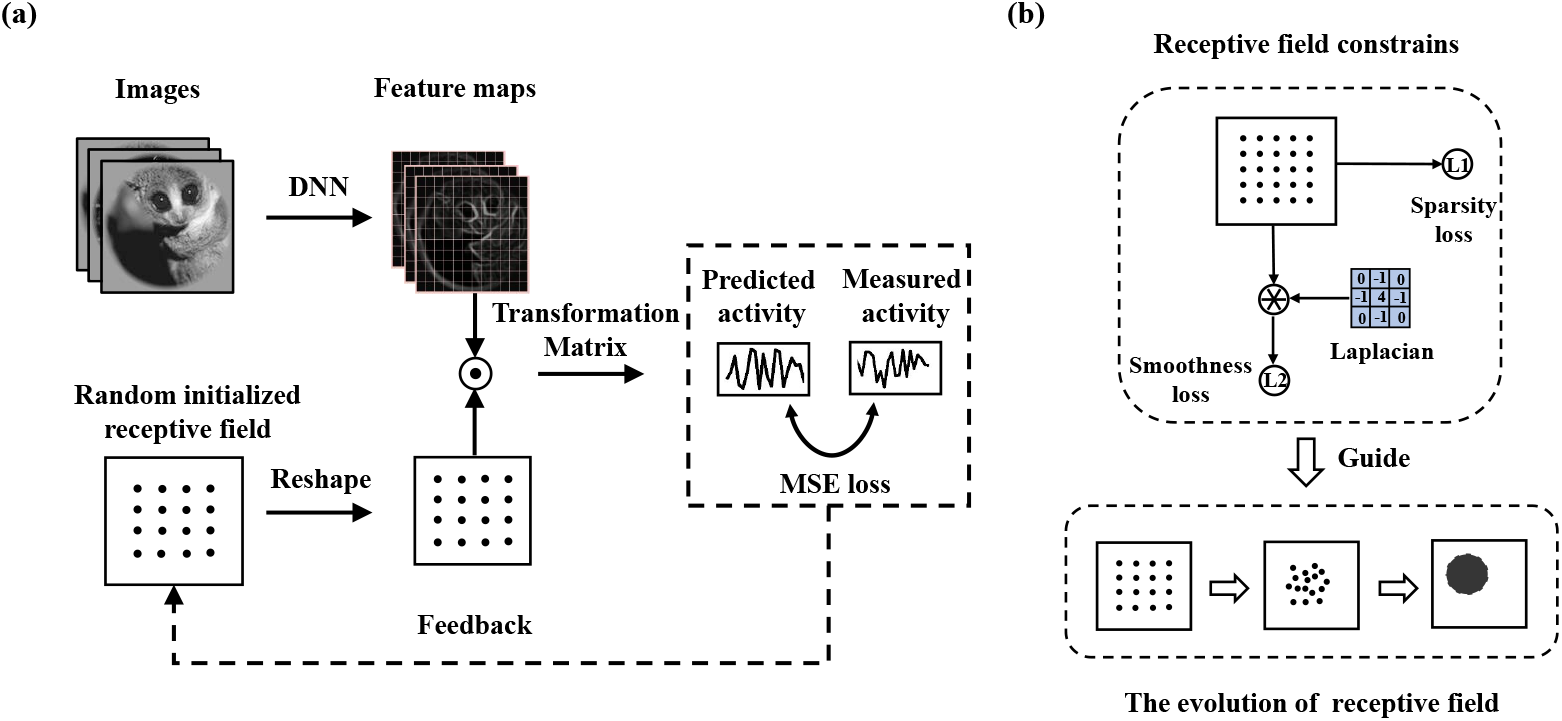
The deatails of the proposed modeling approach. (a) The image data are converted into the feature maps (or attentioned feature maps), and the receptived field is randomly initialized and reshaped to same size as feature maps. In the next phase, the Hadamard product (⊙) of the feature maps and the receptive field produce the refined features. Finally, the refined features simultaneously regressed onto voxel-wise activity by a transformation matrix. (b) The receptive field is regularized by the two forms of regularizations: sparsity and smoothness. These two regularizations are able to guide the evolution of receptive field.

For each specific image **x**_*i*_, its original multi-layer feature maps **h**_*i*_ turn into the refined multi-layer feature vector **r**_*i*_ through the nonlinear feature refinement module. The predicted response at the specific voxel is modeled as a linear combination of multi-layer features. Let us denote the weight of multi-layer features as **w**. In order to predict the response of the specific voxel to the natural image, the weight **w** is element-wise multiplied by the refined feature vector **r**_*i*_. Formally:

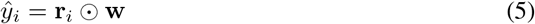

where **r**_*i*_ ∈ ℝ^1×*K*^ and **w** ∈ ℝ^1×*K*^, *K* donotes the number of total feature maps. There is tend to be a voxel-wise bias *b* in practice, and we omit it for notational simplicity, as it does not play a part in the validation accuracy. Finally, the mean-squared error can be formulated as below:

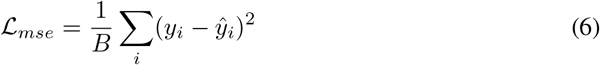

where *y*_*i*_ is the measured voxel-wise activity in response to image *i*, *ŷ*_*i*_ is the predicted activity of the model, B indicates the minibatch size.

### The objective function of the model

The deatails of the whole modeling approach are shown in Fig 4. The final objective function for each specific voxel is defined as follows:

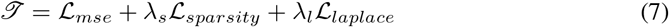

where *λ*_*l*_ and *λ*_*s*_ are hyper-parameters. The first term 𝓛_*mse*_ is the MSE loss. Intuitively, minimizing this loss, which is equivalent to making predicted values approximate true values, can result in more accurate predictions. It is obvious that the L1 regularization term 𝓛_*sparsity*_ plays an sparse role. The RF is supposed to be such a highly sparse area that optimizing the second term contributes to revealing the localized structure of RF, which is beneficial to the interpretability of our model. The third term 𝓛_*laplace*_ is the L2 penalty on the Laplacian of the RF, which is used to make the RF smooth. More Specifically, minimizing this term, the pixels in the particular area of RF tend to be numerically consistent. Therefore, this constraint ensures that RF encodes features within a contiguous region.

When optimizing the objective function, the final goal is to infer RF area and weights of feature maps that lead to more accurate predictions of the voxel’s response. Herein, the objective function can be minimized by Adam Optimizer [36] based on gradient descent method.

## Materials

### Data description

The data uesd in the present study are the public fMRI dataset vim-1 (Data are available at https://crcns.org/data-sets/vc/vim-1.), which are described in detail in [4]. In summary, functional BOLD activity was measured in the occipital lobe with 4T INOVA MR scanner (Varian, Inc.) at a spatial resolution of 2mm × 2mm × 2.5mm and a temporal resolution of 1 Hz. During the acquisition, subjects viewed sequences of 20° × 20° greyscale natural photographs while fixating on a central white square. Photographs were presented for 1s with a delay of 3s between successive photographs.

The data are partitioned into distinct training and testing sets. The training set consists of estimated voxel activity in response to 1750 photographs while the testing set consists of estimated voxel activity in response to 120 photographs. In present study, the fMRI data from visual area V1, V2, V3, V4, LO, V3a, V3b are used for the analysis, and the original training set is further partitioned into training set and validation set. It divides the original data into 5-Fold, and validates each subset (consists of 20% of the original training set) separately, whereas the remaining 4 subsets are used as training set.

### Ethics statement

Two healthy subjects with normal or corrected-to-normal vision participated in the experiments: Subject 1 (male, age 33) and Subject 2 (male, age 25). All subjects provided written informed consent for participation in the experiments, in accordance with the Declaration of Helsinki, and the sharing of vim-1 dataset has been approved by the UC Berkeley Office for Protection of Human Subjects.

### Compared methods

The following models are considered as compared methods:

a. **Compressive spatial summation model (CSS)**: The CSS model takes a contrast image (i.e. an image representing the location of contrast in the visual field) as the input, computes a weighted sum of the contrast image using an isotropic 2D Gaussian, and applies a static power-law nonlinearity [9]. It is an effective way to encode contrast images, while the input pattern limits the flexibility and generalization of the SOC model.
b. **Second-order contrast model (SOC)**: The SOC model starts with a grayscale image (luminance values), applies Gabor filters as a way of computing local contrast, computes second-order contrast within the spatial extent of an isotropic 2D Gaussian, and applies a static power-law nonlinearity [15]. Whereas the CSS model explains only how the location and size of the stimulus relate to the response, the SOC model is more general, explaining the relation of an arbitrary grayscale image to the response. However, its relatively shallow feature representations limits encoding performance, and the progress hits a bottleneck at the natural images.
c. **Feature weighted receptive field (fwRF)**: A latest neural encoding method based on convolutional neural network and multivariate linear regression [13]. It starts with a natural image, obtains feature maps in a pre-trained convolutional neural network, and computes a weighted sum within the spatial extent of an 2D Gaussian receptive field. Finally, it regresses all the feature maps onto brain activity simultaneously and outputs accurate predictions, which outperforms other comparable encoding models to achieve the state-of-the-art performance. However, fwRF only considers the fusion of diverse features from DNNs and still neglects the drawbacks resulted from strong prior assumptions (Gaussian assumption), like previous methods.

### Voxel selection

Voxel selection is a crucial component to fMRI brain encoding, as plenty of voxels may not respond to the visual stimuli in reality. A common approach is to choose those voxels to which the model provided better predictability (encoding performance) during the training process. The goodness of fit between model predictions and measured voxel activities is quantified by the Pearson’s correlation coefficient (PCC). For each voxel, the PCC is computed on the validation set, and is finally an average of 5 runs with different data splits in our experiments. We select voxels with positive PCC for further analyses, and the details of the selected data are summarized in Table 1 (The details of the selected data from subject 2 are shown in S1 Table). Figures in this study refer to data from subject 1 (except Fig 6).

**Table 1.** The details of the selected data from subject 1 in our experiments.

| ROIs | #Instances | #Pixels | #Voxels | #Training | #Validation | #Testing |
| --- | --- | --- | --- | --- | --- | --- |
| V1 | 1870 | 227×227 | 992 | 1400 | 350 | 120 |
| V2 | 1870 | 227×227 | 1529 | 1400 | 350 | 120 |
| V3 | 1870 | 227×227 | 1202 | 1400 | 350 | 120 |
| V4 | 1870 | 227×227 | 1015 | 1400 | 350 | 120 |
| LO | 1870 | 227×227 | 621 | 1400 | 350 | 120 |
| V3a | 1870 | 227×227 | 311 | 1400 | 350 | 120 |
| V3b | 1870 | 227×227 | 210 | 1400 | 350 | 120 |

### Model fitting

The AlexNet [18] architecture pre-trained on ImageNet dataset is exploited to initial both the convolutional and fully-connected layers, and other parameters of the model are randomly initialized. The hyper-parameters of the proposed model are set to (*λ*_*s*_, *λ*_*l*_) = (1, 1), while five-fold cross-validation is carried out to choose better regularization parameters from [0.01, 0.1, 0.5, 1, 5, 10]. In our experiments, the minibatch size *B* is set to 20. The Adam Optimizer [36] with an initial learning rate of 0.00005 and early stopping is adopted. Specifically, we monitor the validation loss every iteration of totally 200 iterations and early stop when the validation loss have not decreased for 5 consecutive times.

### Model Evaluation

To evaluate the encoding performance quantitatively, we use several standard similarity metrics, including **mean squared error** (**MSE**), **Pearson’s correlation coefficient** (**PCC**), and **coefficient of determination** (**COD**, i.e. R^2^). These metrics focus on different properties for the encoding performance. MSE is a common way to evaluate prediction performance in machine learning, which focuses on the point-to-point prediction accuracy. Note that MSE is not highly indicative of predictions, whereas PCC and FEV can take variable texture and goodness of fit into account, which are more significant in neuroscience. We also performed the **statistical significance test (SST)** of model prediction accuracy. For each voxel and each model type, the Pearson’s correlation coefficient between the model prediction and measured response above 0.27 is significant *p* < 0.001 relative to its null hypothesis distribution [4, 13]. In the present study, we use PCC to denotes the prediction accuracy, if there is no special instruction.

### Feature map contribution to the prediction

According to voxel-wise mapping module, all the feature maps contribute linearly to the model prediction. It is a natural way to determine the relative importance of each feature map in terms of the regression weights. In practice, it is difficult to make comparisons across multi-layer feature maps, as the regression weights are dependent on the typical values of each feature map. Hence, in consideration of the linearity of the proposed model, we calculate the Pearson correlation coefficient *ρ*_*l*_ = *cov*(*ŷ, y*)_/_*sqrt*(*var*(*ŷ*)*var*(*y*)) over a subset of feature maps h^*l*^ ∈ h instead of focusing on regression weights. All the disjoint subsets Σ_*l*_h^*l*^ cover all *K* feature maps, and they follows Σ_*lρl*_ = *ρ* where *ρ* is the cumulative PCC between predicted voxel-wise activity and true voxel-wise activity. Here, each *ρ*_*l*_ thereby denotes the contributions of the subset of feature maps to the model prediction.

## Results

### Relationship between CNN layers and brain ROIs

**Fig 5.**
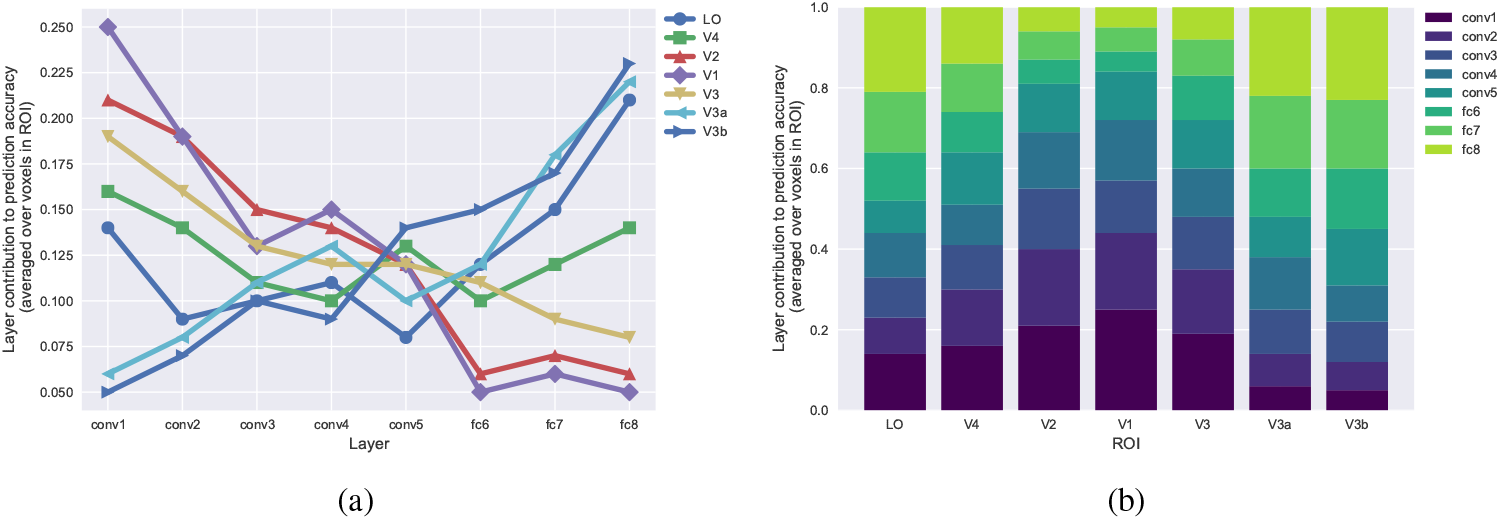
Contributions of the DNN layers to predictions of ROIs. (a) Each column shows the distribution of the single DNN layer contributions to the prediction accuracy of each ROI. The prediction accuracy are averaged over all voxels in each ROI. (b) Each column shows the distribution of each DNN layer contributions to the prediction accuracy for a single ROI. Colored bars within each column indicate the contribution to the prediction accuracy averaged over all voxels in that ROI.

Previous neuroscience studies [21, 24] have shown that the ventral and dorsal visual streams are hierarchically organized, with early visual areas processing low-level visual features (such as edges) and downstream visual areas processing increasingly complex visual features (such as shapes). Does the hierarchical features of CNN have anything to do with the hierarchical visual areas of brain? To answer this question, we analyzed the contributions of different CNN layers to activity prediction in different brain regions-of-interest (ROI). As shown in Fig 5 (a), the contribution of the early visual areas in ventral (V1, V2) and dorsal (V3, V3a, V3b) streams exhibit a clear counter-gradient organization. Contributions of downstream visual areas (V4, LO) are also graded, but are much more uniformly distributed across the CNN layers. Whlie in Fig 5 (b), the contribution of the lowest (conv1) and highest (fc8) layers exhibit a clear counter-gradient organization. Contributions of intermediate DNN layers are also graded, but are much more uniformly distributed across ROIs.

**Fig 6.**
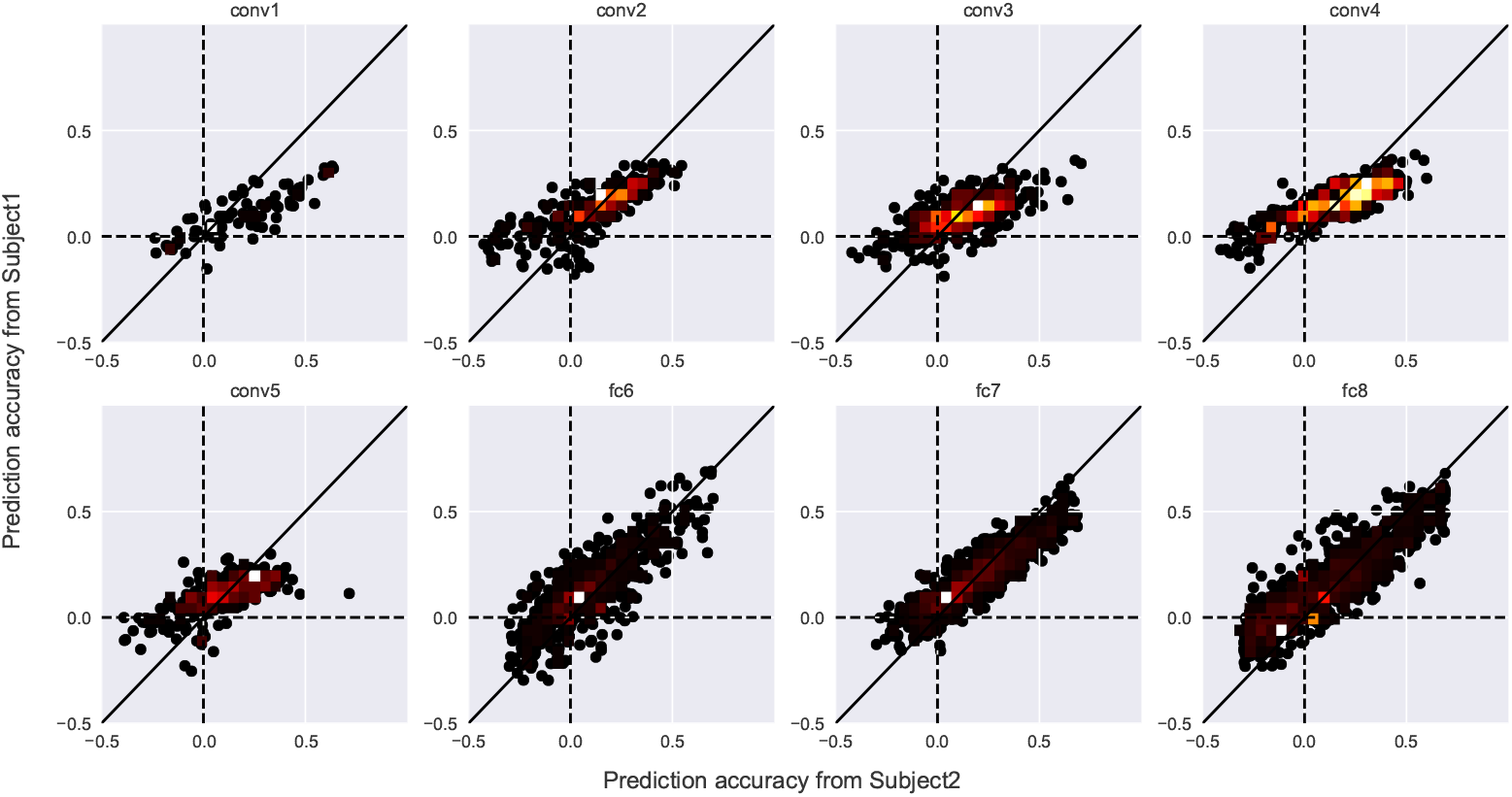
Encoding accuracies of individual feature maps or units (AlexNet) from two subjects of the Vim-1 dataset. Each dot denotes the encoding accuracy of each feature map or unit estimated from Subject1 and Subject2. The color of each dot indicates the density of the plotted dots.

More specifically, the most contributable CNN features for the prediction of the early visual areas come from shallow CNN features, whereas most contributable voxels for the prediction of the downstream visual areas come from the deep CNN features. These results demonstrate a homology between computer and human vision, providing a new opportunity to make use of the hierarchical information from CNN features.

### Individual differences between subjects

To assess the degree of consistency of encodability across subjects, we evaluated the feature map-by-map or unit-by-unit similarity of the prediction accuracy between two subjects of Vim-1 dataset. Fig 6 shows the scatter plots of feature encoding accuracies between Subject1 and Subject2. The prediction accuracies of individual maps or units from the two subjects densely distribute along the diagonal axis for most layers, showing positive correlations between the two subjects. The positive PCCs for all layers of the DNN architectures suggest that the DNN-based neural encoding was highly consistent across subjects even at the feature map or unit level.

### The visualization and convergence of receptive fields

To verify the capacity to estimate receptive fileds of our modeling approach, we intuitively visualized the receptive field with increasing iterations across different ROIs. The results are illustrated in Fig 7, which are the representative voxels from V1, V2, V3, V4, LO. On one hand, it is easy to find that the receptive fields are smooth and localized for the particular voxels. The receptive field shapes may not be regular for all voxels, whereas the main shapes can be clearly distinguished. On the other hand, the preliminary outlines can be formed within 30 iterations while optimization procedure converges within roughly 50 iterations. In practice, for arbitrary voxel, the receptive field can be optimized automatically in this way. The results on the rest of voxels are similar, and we omit them due to space limitations. It can be inferred that, owing to the regularizations, the receptive field in our proposed method is able to capture the reasonable location and extent of pooling over visual features.

**Fig 7.**
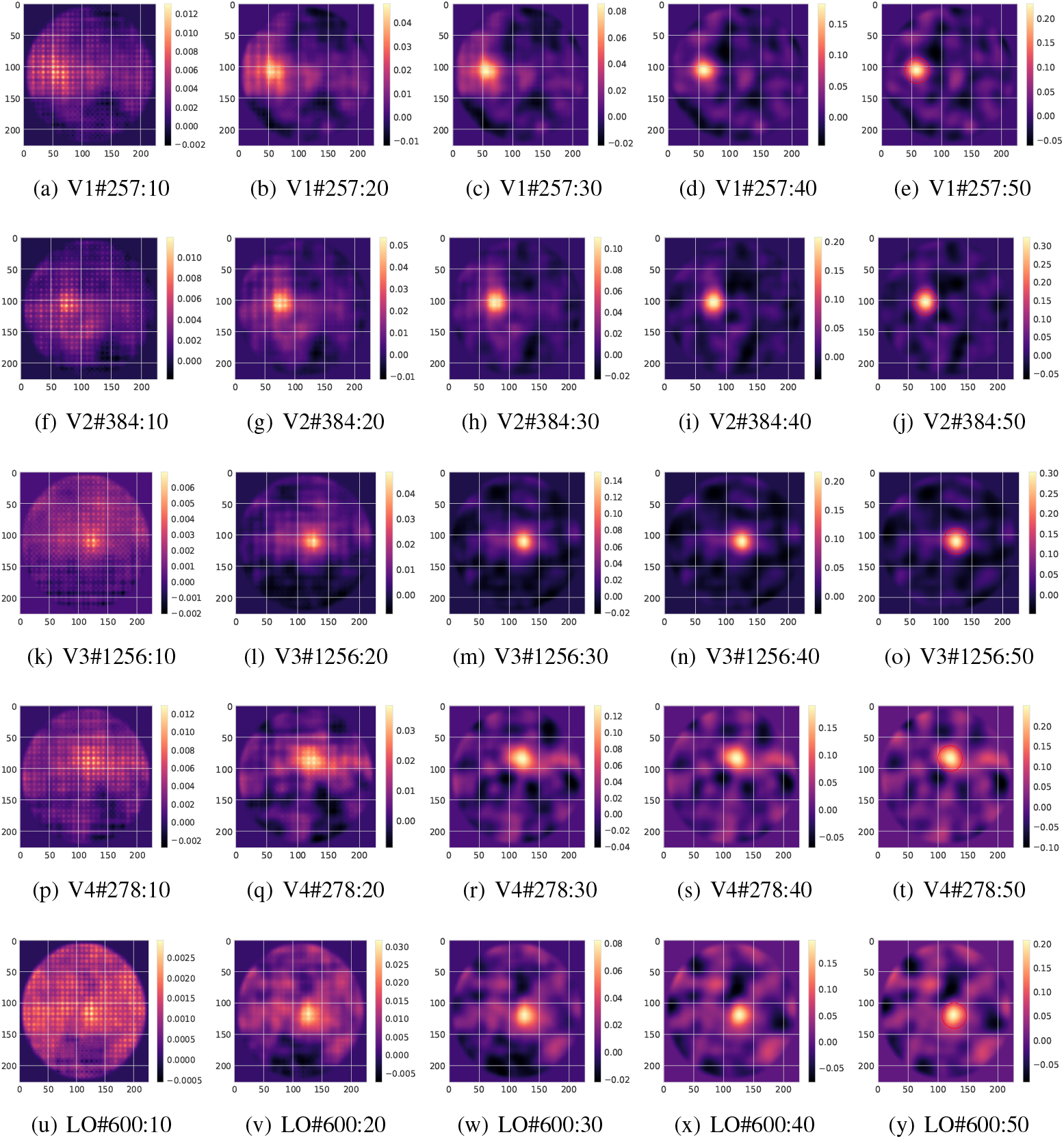
The estimated receptive fileds of the representative voxels with different iterations in ROIs. For example, (a) V1#257:10 shows the results of the 257th voxel in V1 after the 10 iterations. For the particular voxels, the receptive fields are smooth and localized. The preliminary outlines can be formed within 30 iterations while optimization procedure converges within roughly 50 iterations.

### Quantitative analysis of encoding performance

The encoding performce distribution [37] of the proposed method is shown in Fig 8. Voxels located in early visual cortex (V1, V2) are more accurately predicted than those located in higher visual cortex (V4, LO), whileas the difference of encoding performance between most of the visual areas is not very obvious, especially for the MSE metric. It verifies the feasibility of our method in the visual areas. We further compare the proposed method with other baseline methods in terms of three metrics, and the quantitative performance comparisons are shown in Fig 9 and Fig 10 (the details are shown in Table 2). From them, we can find that our method outperforms the baselines in most brain ROIs. Compared with CSS and SOC, the consistently encouraging result shows that the proposed method with a DNN model for visual images is more able to extract nonlinear features from visual images, which may contribute to encoding performance in the primary visual cortex. Furthermore, our method shows obvious better performance than fwRF. In spite of the same feature representation network, it is maybe caused by the fact that fwRF is endowed with the two-dimensional Gaussian assumption and manual parameter space, which may not obtain the global optimal solution of model parameters. In summary, for the current dataset, the substantial superior performance (Consistent results are obtained for subject 2, as shown in S2 Table) of the proposed model verifies that estimating receptive field automatically with weak prior assumptions about the spatial characteristics is beneficial to enhancing the encoding performance.

**Fig 8.**
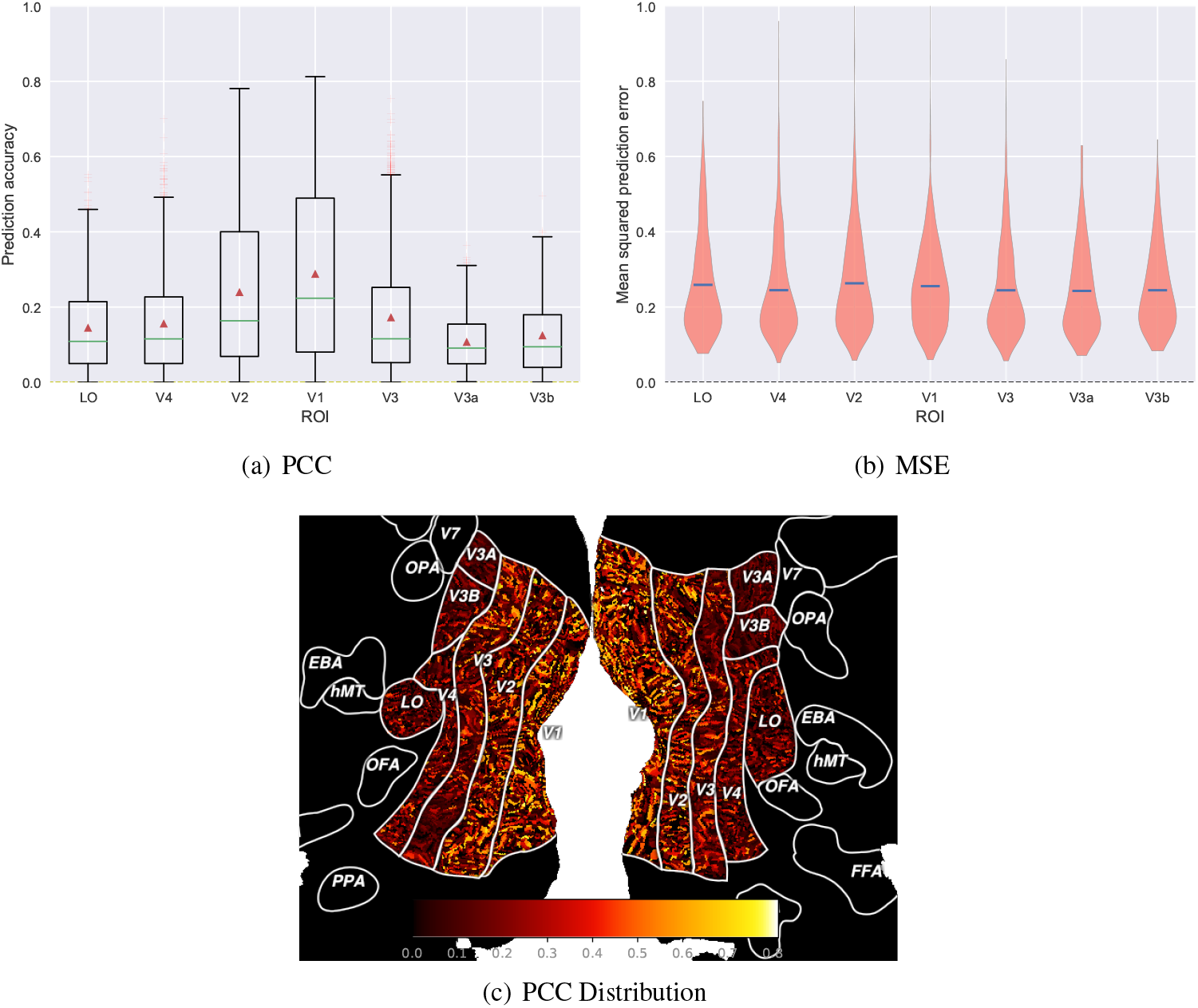
Encoding accuracy for each ROI. (a) Distributions of the encoding accuracy of individual voxels in terms of PCC. Green bars denote median prediction accuracies while red triangles denote mean prediction accuracies averaged across all voxels. (b) Distributions of the encoding accuracy of individual voxels in terms of MSE. Blue bars denote mean prediction accuracies averaged across all voxels. (c) Prediction accuracy of voxels plotted on a digitally flattened map of visual cortex.

**Fig 9.**
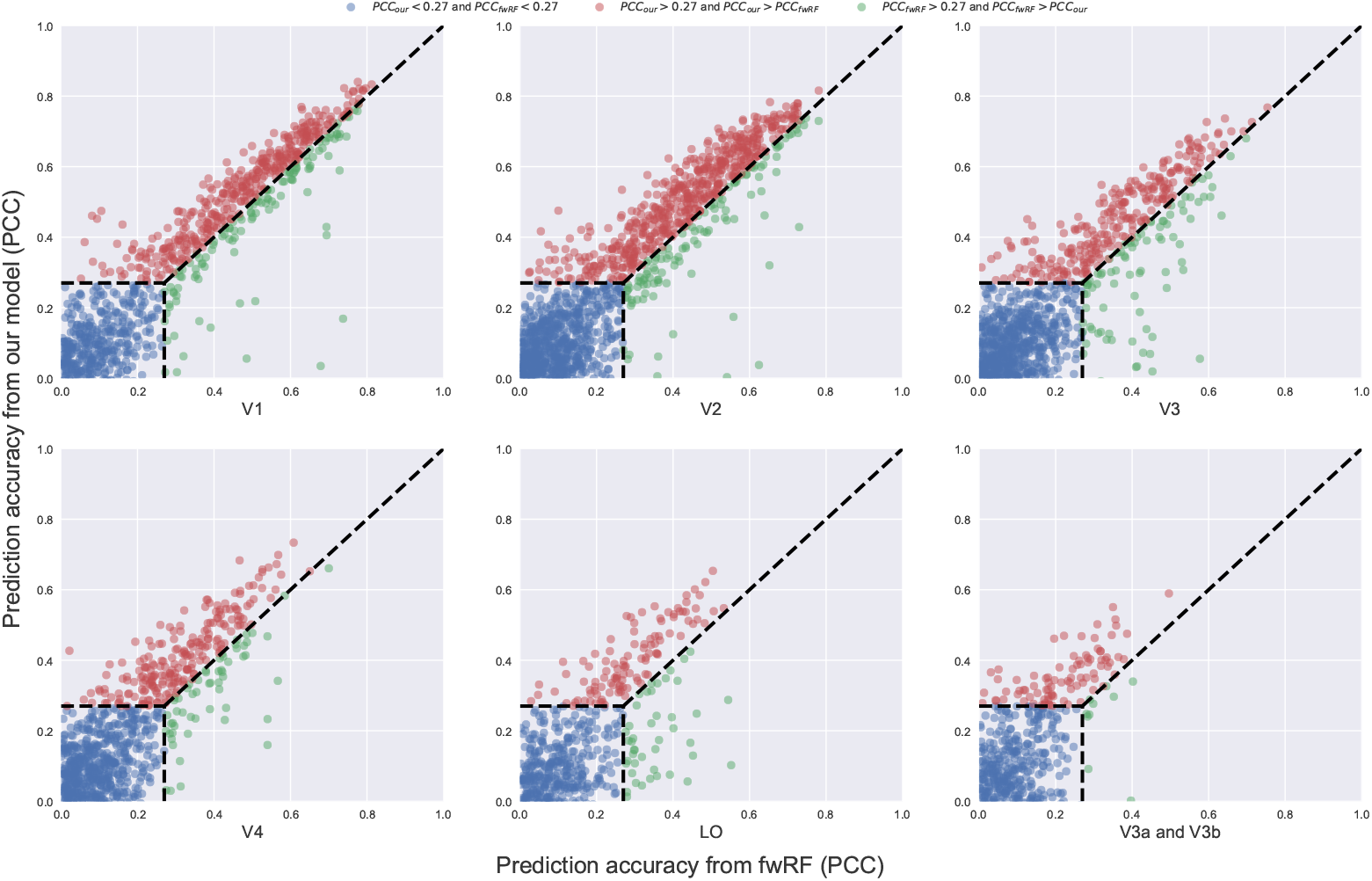
Comparison of prediction accuracy between the proposed method and the fwRF. Each of the six axes displays a comparison between the prediction accuracy of the two models in specific visual ROI (V3a and V3b are plotted in the same axe). In all six scatter plots, the ordinate and abscissa represent the prediction accuracy values of the proposed method and the fwRF. The blue dots indicate the voxels cannot be significantly encoded (under 0.27) by either of the two models. The red dots indicate the voxels that can be better predicted by the proposed method than the fwRF and vice versa for the cyan dots.

**Fig 10.**
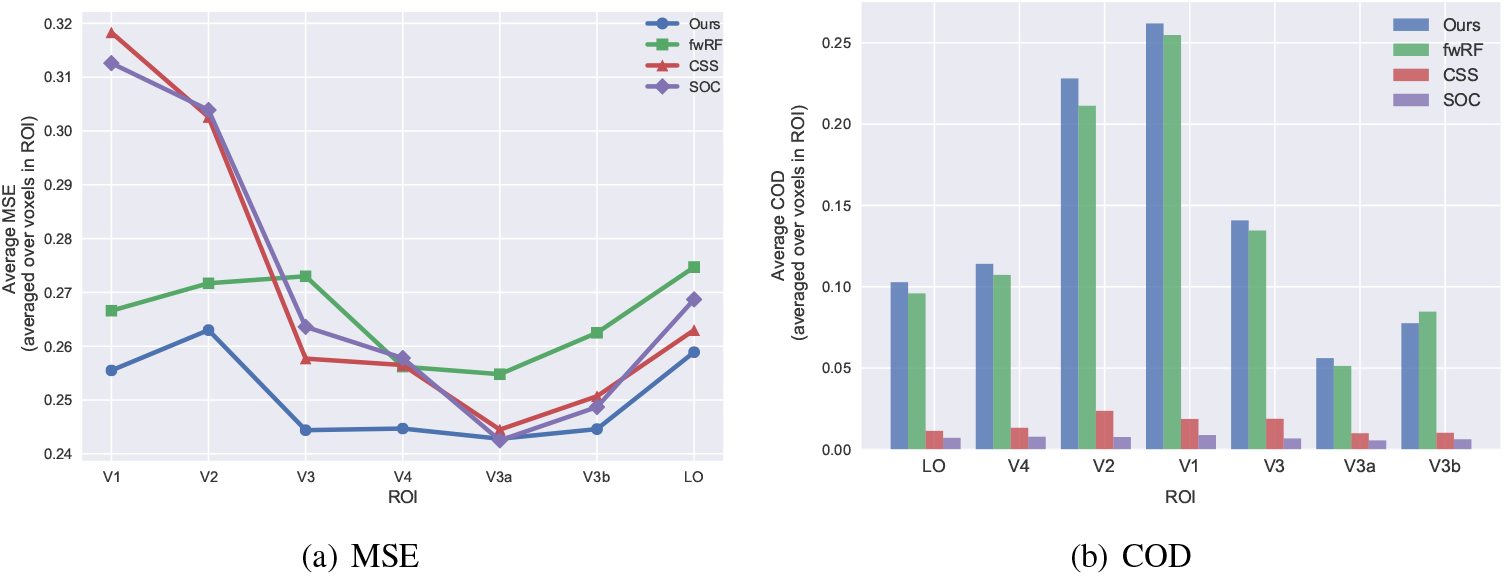
Preformance within different ROIs in terms of MSE and COD. The proposed method outperforms other baseline methods in most brain ROIs

**Table 2.** Performance of several neural encoding models for Subject 1 of the Vim-1 dataset. The best performance (averaged over voxels) within each ROI was highlighted.

| Evaluation | Algorithms | V1 | V2 | V3 | V4 | LO | V3a | V3b |
| --- | --- | --- | --- | --- | --- | --- | --- | --- |
| MSE | CSS | 0.3183 | 0.3026 | 0.2577 | 0.2565 | 0.2630 | 0.2445 | 0.2507 |
|  | SOC | 0.3126 | 0.3039 | 0.2636 | 0.2578 | 0.2687 | 0.2425 | 0.2487 |
|  | fwRF | 0.2666 | 0.2717 | 0.2730 | 0.2562 | 0.2747 | 0.2548 | 0.2625 |
|  | Ours | **0.2555** | **0.2630** | **0.2444** | **0.2447** | **0.2589** | **0.2428** | **0.2446** |
| PCC | CSS | 0.0448 | 0.0463 | 0.0330 | 0.0321 | 0.0221 | 0.0261 | 0.0222 |
|  | SOC | 0.0911 | 0.0611 | 0.0483 | 0.0496 | 0.0535 | 0.0171 | 0.03041 |
|  | fwRF | 0.2886 | 0.2400 | 0.1600 | 0.1517 | 0.1257 | **0.1097** | 0.1251 |
|  | Ours | **0.2943** | **0.2468** | **0.1730** | **0.1565** | **0.1453** | 0.1077 | **0.1363** |
| COD | CSS | 0.0187 | 0.0237 | 0.0188 | 0.0133 | 0.0114 | 0.0099 | 0.0102 |
|  | SOC | 0.0088 | 0.0076 | 0.0067 | 0.0078 | 0.0071 | 0.0055 | 0.0062 |
|  | fwRF | 0.2548 | 0.2113 | 0.1346 | 0.1073 | 0.0959 | 0.0513 | **0.0847** |
|  | Ours | **0.2619** | **0.2281** | **0.1408** | **0.1141** | **0.1028** | **0.0561** | 0.0776 |

### Sensitivity analysis

The sensitivity analysis is crucial to DNN-based models. Since weak prior assumptions (sparsity and smoothness regularizations) are adopted in our model, we also performed the sensitivity analysis. It involves two hyper-parameters Θ = (*λ*_*s*_, *λ*_*l*_), which need setting properly. For the sake of studying the sensitivity of the model with respect to different values of these parameters, we plot effective encoding results (average over voxels with the prediction accuracy higher than 0.5) with different values of regularization parameters, and displayed the results in terms of PCC and MSE in Fig 11. The figures for variation of MSE and PCC show the same pattern, and model performs stably with the variation of different *λ*_*s*_ and *λ*_*l*_. The results also suggests that the best regularization parameters *λ*_*l*_ and *λ*_*s*_ can be chosen from [0.5, 1, 5] and [1,5], where the proposed model achieves good results.

**Fig 11.**
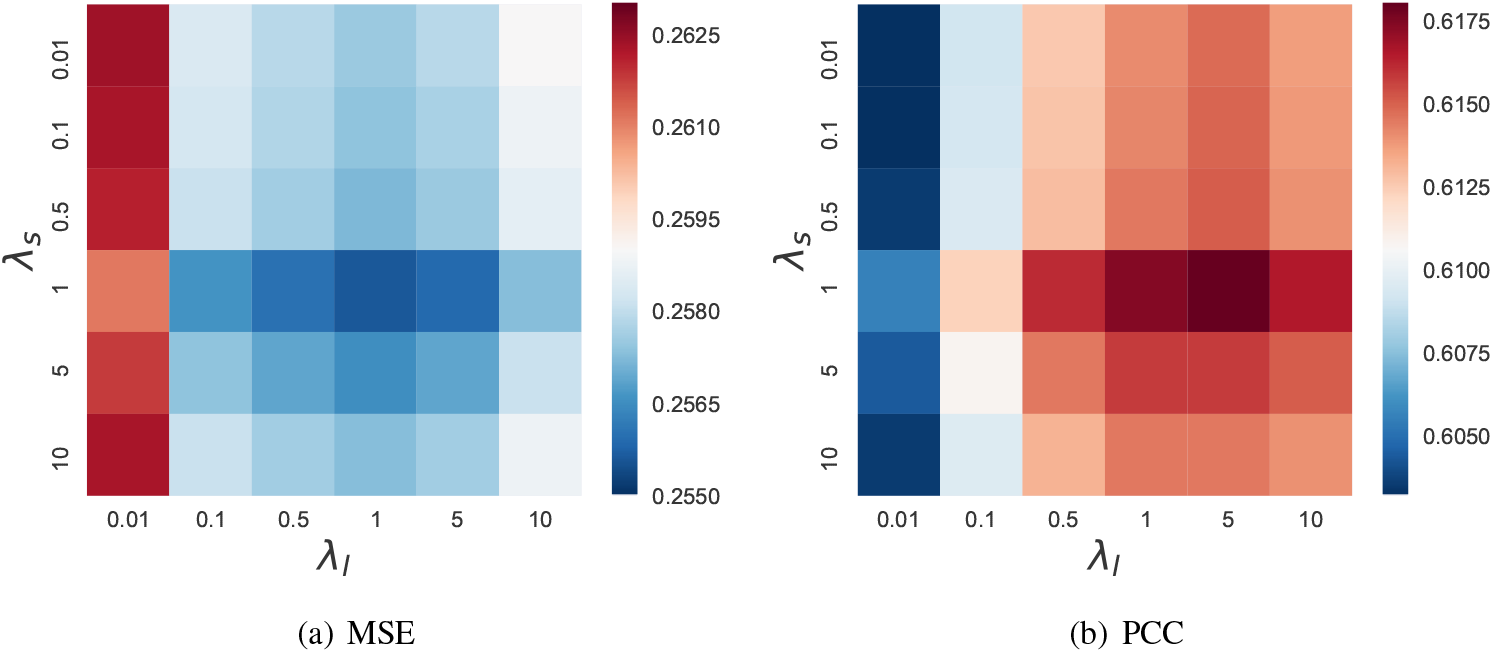
Encoding perfomance with the variation of *λ*_*l*_ and *λ*_*s*_ in terms of MSE and PCC. The best regularization parameter *λ*_*l*_ and *λ*_*s*_ can be chosen from [0.5, 1, 5] and [1,5], where the proposed model achieves good results.

### Extension for multi-voxel joint encoding

In the visual cortex, the responses to stimulation in the classical receptive fields can be modulated by stimulation in the extra-classical receptive field [7]. These modulations can be excitatory or inhibitory and have been characterized in detail by electrophysiological and psychophysical studies [38–40]. It inspired us to relate receptive fields of multiple voxels to mimic this modulations and construct multi-voxel joint encoding models. Here, we use the similarity loss to relate the adjecent voxels, as voxels in the same area tend to perform similar computations. Taking two-voxel cooperative encoding as an example, for voxel *j* and voxel *k*, to ensure their receptive fields as similar as possible, we use an L2 penalty on the difference between receptive fields with the strength *λ*_*m*_:

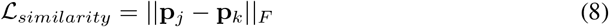

The joint objective function of the multi-voxel joint encoding model is formulated as follow:

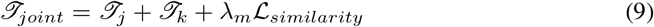

Here, *ℐ*_*j*_ and *ℐ*_*k*_ are the objective function of voxel *j* and voxel *k*, respectively, and 𝓛_*similarity*_is the bond between two voxels. This bond can be extended to three or more voxels according to appropriate definitions of voxel neighborhood. In the present study, we did a preliminary verification of the multi-voxel joint encoding.

In consideration of computational similarities in the same visual area, we define a voxel neighborhood that consists of three voxels according to their location index in the Vim-1 dataset. Those voxels (up to three voxels) whose location index are adjacent are chosen to be jointly optimized. The hyper-parameter *λ*_*similarity*_ is set to 1, while five-fold cross-validation is carried out to choose the better regularization parameter from [0.1, 0.5, 1, 5]. We compare the multi-voxel joint encoding with the single-voxel encoding, and the results are shown in Fig 12. It can be found that, both in early visual areas (V1, V2, V3) and higher visual areas (V4, V3a, V3b and LO), after the multi-voxel joint encoding, a slight shift of the voxels toward the right indicates an advantage for the multi-voxel joint encoding. The results suggest that, owing to voxel neighborhood information, the multi-voxel joint encoding is conducive to rescuing voxels with poor signal-to-noise characteristics.

**Fig 12.**
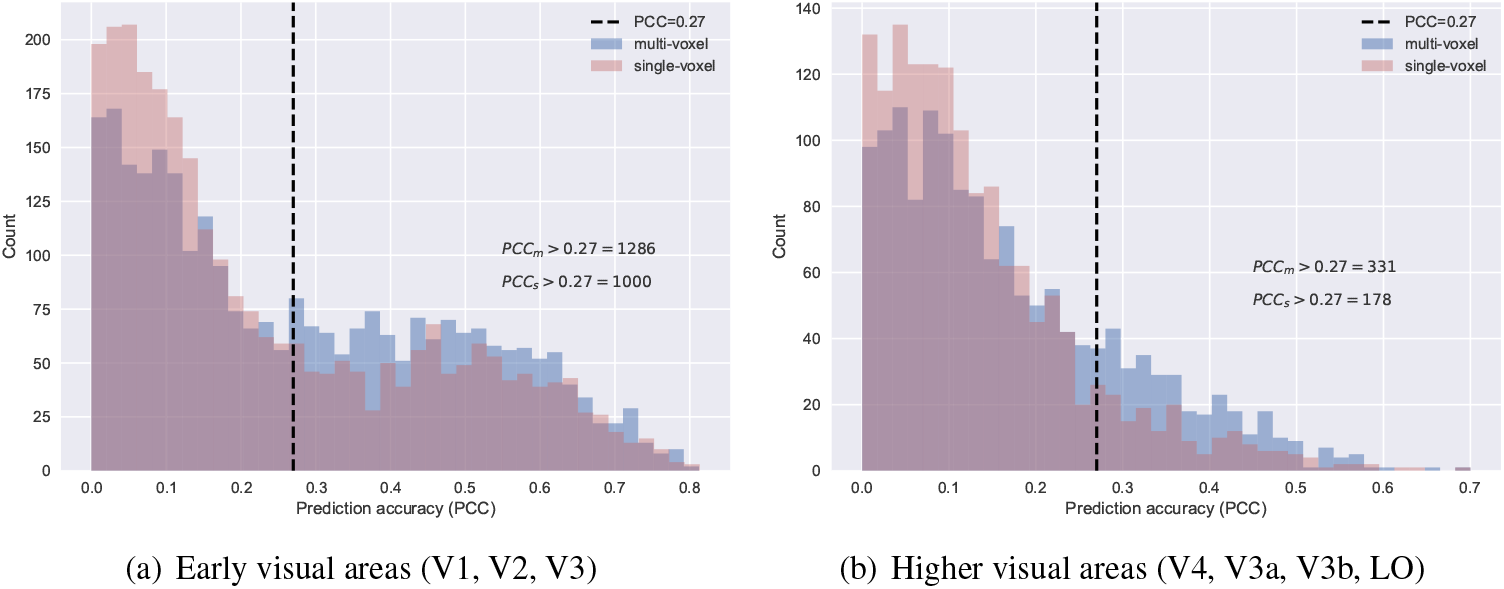
Comparisons between the single-voxel encoding and the multi-voxel joint encoding. (a) Histogram of voxels within early visual areas (V1,V2,V3). There are 1286 significant voxels in the multi-voxel joint encoding model, in contrast to 1000 significant voxels in the single-voxel encoding model. (b) Histogram of voxels within higher visual areas (V4,V3a,V3b,LO). The number of significant voxels in the multi-voxel joint encoding is 331 while that in the single-voxel encoding model is 178. Results in both early visual areas and higher visual areas demonstrate that the multi-voxel joint encoding is conducive to rescuing voxels with poor signal-to-noise characteristics

## Discussion

The proposed model is a new approach to building voxel-wise neural encoding models. Attributing to the nonlinear feature refinement and separability of “what” and “where” parameters, the proposed model is endowed with explicit receptive fields and feature maps for each voxel, which facilitates the interpretability of neural encoding models. Compared with either traditional or deep learning methods, the proposed model achieved the superior performance in visual brain areas.

### Relationship to previous work

Preceding the proposed modeling approach, a number of important voxel-wise modeling approaches have made significant progress in neural encoding for human visual cortex. One important class of models are those designed specifically for retinotopic modeling. This class involves the inverse retinotopy method [41] and the population receptive field methods [6–8]. These methods implement estimation and analysis of the locations and sizes of voxel-wise receptive fields, with the dedicated retinotopic mapping experiments that utilize the artificially designed stimuli. However, they do not provide explicit feature maps and turning functions directly and require fussy in-silico experiments. Another effective class of models are those designed based on DNNs [13]. These methods are special cases of the general linearized regression approach, they construct a set of nonlinear features from deep neural networks and present explicit feature map. Nevertheless, they still limit the receptive field with Gaussian assumptions in advance. The strong prior assumptions about receptive field have the advantage of reducing the number of parameters of the model, whereas they reduces their expressiveness as well.

The proposed modeling approach with weak prior assumptions about receptive fields is a more special case of the general linearized regression approach, and it also depends on the construction of a set of nonlinear features that result from deep neural networks. This modeling approach overcomes two limitations of previous general regression approaches. First, in general regression approaches, the shape of receptive field is pre-defined, which makes the model prone to erroneous eatimation of receptive field characteristics. Second, under the general approach deriving an explicit receptive field and feature tuning function often requires grid searching from plenty of candidate receptive fields. However, it is an kind of effective but not sensible way to set search parameters (such as total number and the search interval of candidate receptive fields) according to experience, as it demands too much manual effort.

### Advantages of the proposed model

There are at least three possible reasons for why the proposed model outperforms the other compared methods. One primary possible reason is the flexible receptive field of the proposed modeling approach. Structure analysis of receptive fields and preformance comparisons with fwRF suggest that the model takes advantage of this flexibility by the weak assumptions about spatial characteristic of rereceptive fields. Without defining the shape of receptive field in advance, the model can make full use of the distribution of training data and estimate the optimal receptive field automatically, revealing the efficient location and extend of visual features. Another possible reason is the usage of channel attention module. On account of it, our model is able to select inter-layer feature attributes and reducing redundant information. Furthermore, strengthening important ones and weakening unimportant ones are conducive to subsequent cross-layer regression. The third possibility is the space-feature separability derived from the proposed model. Explicit receptive fields in the nonlinear feature refinement module are able to capture spatial information, whereas explicit weights of feature maps allow the proposed model to select cross-layer combinations of visual features that closely resemble the feature selectivity of each voxel. We infer that the increased expressiveness is made possible by this space-feature separability.

### Limitations of the proposed model

While the proposed modeling approach exhibits good encoding performance, there is still an imperfection worthy of consideration: the measurement of receptive field size. The proposed modeling approach yields an explicit receptive fields with weak prior assumptions, which provides pros and cons. It facilitates the expressiveness of model, whereas brings the difficulties to the measurement of receptive field size. The receptive field size is different from classical population receptive field (pRF) [6]. The differences underline the fact that the eatimation of receptive field in our method depends entirely upon the feature maps. As the proposed modeling approach does not impose an strong assumptions (regular shapes) on receptive fileds, it limits the measurement of receptive field size to a certain esxtent. However, for irregular receptive field, we can supply an alterbnative measurement method, which is the blob detection. In light of different pixels of the receptive field, blob detection can detect an approximate regular shape, such as the Gaussian shape (areas circled by red line in Fig 7 (e)).

### The receptive field structure

The receptive field presented here is a mask with sparsity and smoothness regularizations, and the salient characteristic is that its shapes can be irregular. While keeping the shape regular has the advantage of reducing model parameters of the model, it also reduces the expressiveness. In particular, the other appropriate operators instead of Laplacian operator can be adopted in our method, exploring more appropriate receptive field structures. For example, the Laplacian of Gaussian operator may allow the model to explicitly capture receptive fields with a “Mexican hat” profile that enforce a suppressive band around an excitatory center. Furthermore, the receptive field with any appropriate weak prior assumptions could be trained using the proposed modeling approach presented here. There may be a more optimal assumption about receptive field structure, which needs future studies to confirm.

### Encoding in higher visual areas (V4 and LO)

It is worth noting that all the presented methods obtain good performance in early visual areas (V1,V2 and V3) while little effect on higher visual areas (V4 and LO). Nevertheless, the proposed modeling approach is still superior to other methods, which makes progress in higher visual areas. However, it is meaningful to build an effective encoding model in higher visual areas, as it may reveal the high-level visual information processing in the visual cortex. Deep neural network has mapped the function of the human visual cortex [35] and revealed a gradient in the complexity of neural representations across the ventral stream [21]. The convolutional layer encoded location and orientation-selective features while the fully-connected layer encoded semantic features. In order to make progress in higher visual areas, perhaps more attention to the fully connected layer will make sense. The receptive fields easimated in our model focus on the convolutional layer, whereas the fully-connected layer are neglected. It is worth our while further improving the encoding performance of our proposed modeling approach with appropriate operations on the fully-connected layer.

## Supporting information

**S1 Table. The details of the selected data from subject 2.**

**S2 Table. The quantitative results of encoding performance for Subject 2.**

## Acknowledgments

This work was supported in part by the National Key Research and Development Program of China (2018YFC2001302), in part by National Natural Science Foundation of China 91520202, 61602449, in part by Chinese Academy of Sciences Scientific Equipment Development Project (YJKYYQ20170050), in part by Beijing Municipal Science and Technology Commission (Z181100008918010), in part by Youth Innovation Promotion Association of Chinese Academy of Sciences, and in part by Strategic Priority Research Program of Chinese Academy of Sciences (XDB32040200).

